# Ontogeny and functional potential of founding dendritic cells in the developing lung

**DOI:** 10.1101/2025.09.30.679620

**Authors:** Radika Soysa, Saajidah Abideen, Victor Zepeda Reyes, Mark B. Headley

## Abstract

Lung development begins in utero and reaches full maturity post birth. Dendritic cells (DC) play a key role in immune regulation in lungs. However, comprehensive exploration of DCs in these immature lungs has not been performed. Here we explored DCs from fetal to newborn mouse lungs phenotypically, ontogenetically, transcriptomically and functionally and found two DC subsets, resembling adult cDC1 and cDC2, but with key differences. Phenotypically, fetal-cDC1 lacks the classical-DC1 (cDC1) marker XCR1, while the fetal-cDC2 express both cDC-associated genes as well as monocyte-derived DC genes. Both DC subsets wane as lungs enter the alveolar stage, giving way to the more familiar adult cDC1 and cDC2. Both fetal-cDC1 and fetal-cDC2 derive from ED14.5 fetal liver Macrophage Dendritic Progenitors, not from monocytes or classic Precursor-cDC (Pre-cDC), indicating a unique ontogeny of first DCs in developing mouse lungs. Together we provide the first in depth exploration of first DCs in developing lungs.

## Introduction

At birth, human and mouse fetuses experience a major environmental shift, leaving the protective *in utero* environment and entering the outside world. Accompanying this process, lungs undergo a mechanical transformation, for the first time, cycling between deflated and inflated states that are essential for respiration and the survival of the newborn. In mice and humans, lung development begins in early fetal stages and then proceeds through a tightly orchestrated differentiation process to increase in cellularity and complexity (Schittny, 2017).

Morphologically, lung development is divided into 5 stages, namely embryonic, pseudoglandular, canalicular, saccular and alveolar stages (Schittny, 2017). At birth, mouse lungs exist in the saccular developmental stage, identified by simplified air sacs and developed vasculature, equipped to support gas exchange (Loering et al., 2019; Warburton et al., 2010). Post birth, at day 5 lungs progressively transition into the alveolar stage where the process of alveolarization dramatically increases the surface area of the lungs to maximize gas exchange efficiency. In humans, term-born infant lungs are in the early alveolar developmental stage, while lungs of pre-term infants are still within the saccular developmental stage (Hilgendorff et al., 2014). In this transition lungs endure substantial changes in oxygen tension accompanying the transition from the *in utero* environment at 30-50 mmHg pO_2_ to the external environment at 90-100 mmHg pO_2_ (Gebb and Jones, 2003; Vogel et al., 2015). Furthermore, at birth, lungs begin to encounter airborne microbes and particles, including harmful antigens that can trigger inflammatory responses. If not regulated these inflammatory responses can be detrimental to the survival of the newborn. In part due to the under-developed state of the lungs and susceptibility to inflammation-induced damage, pre-term infants are at a high risk for developing chronic lung inflammation leading to Bronchopulmonary dysplasia (BPD) a debilitating chronic disease that negatively impacts respiration, susceptibility to infection, and overall well-being (Davidson and Berkelhamer, 2017). Thus, a well-regulated immune response in the lungs is essential for the overall health of the newborn.

Dendritic cells in the lungs are essential for the surveillance of antigens and initiation of adaptive responses. In adult steady-state lungs, there are two major types of classical dendritic cells (cDC) –cDC1 and cDC2. Both subsets respond to inflammatory stimuli, secrete cytokines, and regulate the activities of downstream effector cells, for example T cells and Natural Killer (NK) cells (Cook and MacDonald, 2016; Lambrecht and Hammad, 2012). While rare at steady-state, in inflammatory conditions monocyte-derived DCs (moDC) populate lungs and take part in adaptive responses (Guilliams et al., 2013b). A small number of studies have profiled the lung DC compartment in the early stages of life (Cui et al., 2023; Cui et al., 2021; Ruckwardt et al., 2018). Phenotypic analyses done by de Kleer et.al. of the mouse lung DC compartment between embryonic day (ED) 20 and postnatal day (P) 21 suggested that cDC1s defined as CD11c^+^, MHCII^+^, CD11b^-^ cells are exceedingly rare in saccular stage while moDCs are the major DC type until CD11b^+^ cDC2 take over in the early alveolar developmental stage (de Kleer et al., 2016). Another study by Silva-sanchez et.al. identified two cDC1 subsets in post-birth lungs based on expression levels of CD103 antigen suggesting that phenotypically distinct cDC1s are present in early-life lungs (Silva-Sanchez et al., 2023). In addition, a study done on human fetal tissues indicate that lungs of pseudoglandular and canalicular stages contain cDC1 and cDC2 supporting that DCs seed developing lungs in fetal stages (McGovern et al., 2017).

However, no studies have carefully examined the saccular stage lungs, corresponding to human preterm lungs and newborn murine lungs, for the presence of cDCs and an analysis of the progressive changes in diversity, phenotype and functions of fetal lung DC within the saccular stage has not been performed. Studies that modeled the lung diseases BPD (Cui et al., 2021), allergy (de Kleer et al., 2016) and viral (Malloy et al., 2017; Ruckwardt et al., 2014; Ruckwardt et al., 2018) infection in neonatal mice have suggested that early alveolar stage CD103 DCs respond distinctly to their adult counterparts, however, functional analysis of saccular stage DCs has not been performed.

Here, we present an extensive phenotypic and transcriptomic analysis of the early-life lung DC compartment within fetal and neonatal development, encompassing canalicular, saccular and early alveolar lung development stages. We have identified a major population of cDC1s, which we termed fetal-cDC1 that co-exist with a cDC2-like population of DCs previously termed moDC based on their expression of CD64 (de Kleer et al., 2016), a marker of monocyte-derived cells. Both fetal-cDC1 and cDC2-like cells seed lungs in the canalicular stage, remain through saccular stage and wane as lungs enter the alveolar stage. Functional investigations of these fetal-cDC1 and cDC2-like cells indicated that they are mature DCs with a capacity for activation, cytokine production upon stimulation, and possess the ability for antigen uptake, processing and presentation to T cells. Ontogenetic analysis of these fetal-cDC1 and cDC2-like DCs using in vitro strategies revealed that ED14.5 fetal liver Macrophage Dendritic Progenitors (MDP), in contrast to fetal liver monocytes or adult bone marrow derived MDPs, can give rise to both subsets suggesting that these saccular stage DCs have a distinct fetal ontogeny compared to adult DCs, and support a re-classification of the early life cDC2-like cells from moDC to fetal-cDC2. Here we provide the first in depth analysis of saccular stage lung DC.

## Results

### Saccular developmental stage of lungs contains phenotypically distinct dendritic cells from alveolar developmental stage

When mice are born their lungs are in the saccular developmental stage. Around day 5 post-birth lungs begin transitioning into the alveolar developmental stage. From here onwards lungs undergo alveolarization and give rise to fully mature lungs by the fifth week of life (Fig. 1 A). To explore dendritic cells (DC) in lungs at birth we compared newborn lungs (P1) to 8-weeks old adult lungs using Vectra multiplex immunofluorescent microscopy (mIF). Morphologically, as expected, P1 lungs showed simplified alveoli with emerging vasculature and airways and had thicker regions of parenchyma (Fig. 1B). In contrast, adult lungs were fully alveolarized with well-developed airways and vasculature connected via thin regions of parenchymal tissue. To identify DCs in these lungs we performed mIF for Class II major histocompatibility complex molecules (MHCII) and integrin alpha E (CD103) antigens which are known to be expressed on neonatal lung DC (Lambrecht and Hammad, 2012). We found MHCII^hi^ cells that are dispersed in parenchyma in P1 lungs and among them a subset expressed CD103 (Fig. 1B). To further describe these MHCII^hi^ cells we employed flow cytometry. We developed a panel to identify DC in lungs based on current literature (Cabeza-Cabrerizo et al., 2021). Using this panel, we examined the lung DC compartment in mice from embryonic day (ED)16.5 to 9 days of age and contrasted to 8-week-old adult lungs. This range includes canalicular, saccular or alveolar developmental stages (Fig. S1A). After gating on live single cells, we defined antigen presenting cells as Lineage negative (CD3, CD19, B220, Ly6G, NK-1.1, Nkp46, SiglecF, Ter119), CD45^+^CD11c^+^MHCII^+^ cells. Within this population we enumerated cDC1 as XCR1^+^CD24^+^ cells and cDC2 as SIRPa^+^CD11b^hi^CD64^-^ cells. Cells that expressed CD64 within the SIRPa and CD11b^hi^ population were identified as moDC-like cells based on prior reports (de Kleer et al., 2016). We found that fetal lungs as early as ED16.5 contained DCs (Fig. S1B). Interestingly, DCs in fetal lungs i.e canalicular and early saccular development stages prior to birth did not contain cDC1 or cDC2 per current definition of these subsets (Fig. 1C and Fig. S1B). Instead, these early lungs had a major population of moDC-like cells, consistent with previous reports (de Kleer et al., 2016). Further, and in contrast to adult tissues, fetal lungs contained a large subset of CD11c^+^MHCII^+^ cells lacking expression of either the canonical cDC1 marker XCR1 or the cDC2 marker SIRPa, however, otherwise phenotypically resemble cDC1, expressing CD103 and CD24 within the context of our flow cytometric panel (Fig. 1C and Fig. S1D and E).

**Fig. 1.**
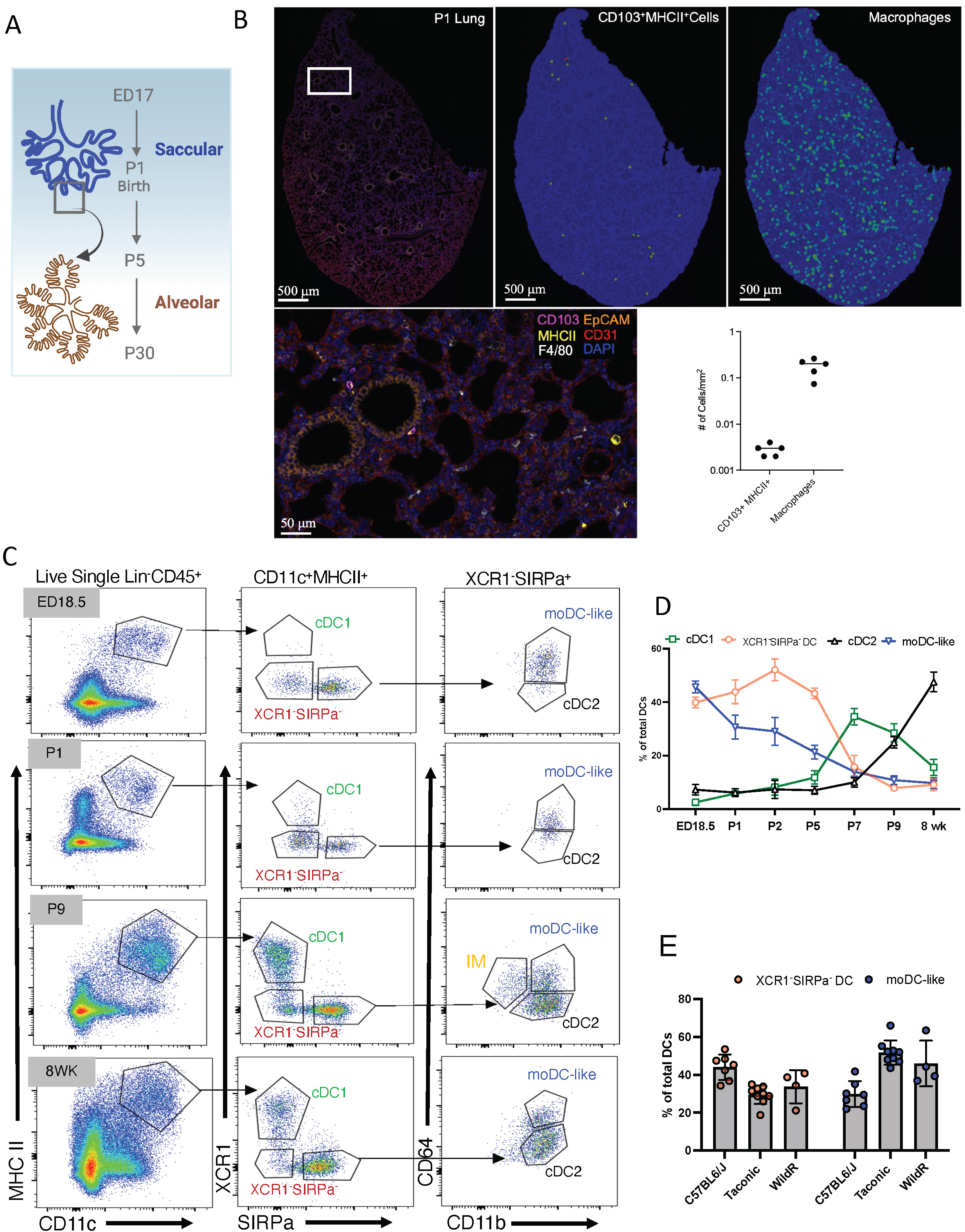
Dendritic cells in saccular developmental stage are phenotypically different from alveolar stage. (A) Illustration of lung developmental stages in fetal to neonatal transition. ED: Embryonic Day, P: Postnatal (B) Representative immunofluorescence images indicating antigen presenting cells in P1 lungs. Bar graph indicates the CD103^+^MHCII^+^ DC and macrophages per area. Two independent experiments performed with n>3 per group. (C) Representative FACS plots indicating the developing DC compartment (gated on Live Lineage (CD3 B220 CD19 NK1.1 SiglecF NKp46 Ter119)^-^CD45^+^CD11C^+^ MHCII^+^ cells) at ED18.5 and post-natal P1, P9 and 8-week time points. cDC: conventional DC, moDC-like: Monocyte-derived DC-like, XCR1^-^ SIRPa^-^DC and IM: interstitial macrophages (D) Time course analysis of frequencies (mean ± SD) of each DC subset within total DCs during lung development from ED18.5 to 8 Weeks. (E) Comparison of percent XCR1^-^ SIRPa^-^DCs and moDC-like cells at P1 in total DCs among 3 colonies of C57/BL6 mice; house bred mice from Jackson(C57BL6/J), mice from Taconic and mice with wild commensal flora (WildR). C, D and E experiments were performed two times with n≥4 group.

To explore the relative proportions of these putative early-life DC subsets (cDC1, cDC2, XCR1^-^SIRPa^-^ and moDC-like cells), during lung development, we analyzed the frequency of each subset from ED16.5 (Fig. S1 B) through ED18.5 to adult (8-10 weeks of age) (Fig. 1D). FACS analysis indicated that in developing lungs, moDC-like cells and this previously undescribed XCR1^-^SIRPa^-^ subset are present as early as ED16.5, corresponding to early canalicular stage and remained the major CD11c^+^MHCII^+^ populations until the end-saccular stage around post-natal day 5. At birth (P1), XCR1^+^ cDC1s first appear as a rare population and increase in frequency as lungs transition into alveolar stage consistent with prior reports (de Kleer et al., 2016). Similarly, cDC2 began to appear post-birth and became the major subset of DCs in the adult lungs. Towards the end of the saccular developmental stage, CD11c^+^MHCII^+^ interstitial macrophages (IM) defined as CD11b^lo^CD64^+^Mertk^+^ began to appear (Fig. S1F), while these cells were not present in lungs during fetal stages and at birth (Fig. 1C). Next, we compared the newborn lung DC compartment in our inhouse-bred C57/BL6 strain (originally derived from Jackson Labs) to C57/BL6 mice from Taconic, and C57/BL6 mice with wild commensal flora (WildR) and found that moDC-like cells and XCR1^-^SIRPa^-^ DCs remained the major subsets of DCs at birth while cDC1s remained a rare population (Fig. 1E). These data suggest that the phenotypic differences in DCs in developing lungs are conserved in mice from different backgrounds and with diverse flora.

Together, our data so far replicated previous reports that moDC-like cells are a major DC subset in the neonatal period that wanes in frequency as lungs mature. In addition, we found an equally prominent lung DC population, phenotypically defined as Lin^-^ CD45^+^ CD11c^+^ MHCII^+^ XCR1^-^ SIRPa^-^ in the fetal and newborn lungs that decreased in frequency as lungs transitioned into alveolar developmental stage. Conventionally defined cDC1s and cDC2s increased in frequency post-birth and became the prominent lung DC subsets as lungs transition into alveolar stage, also as previously described.

### XCR1^-^SIRPa^-^ DCs display key features of cDC1

Next, we examined the relationship between XCR1^-^SIRPa^-^ DCs, moDC-like cells, and conventionally defined cDC1 and cDC2. For this we first sorted cDC1, XCR1^-^SIRPa^-^ DC and moDC-like cells from P1 lungs, and cDC1, cDC2 and IM from P9 lungs, and performed bulk-RNAseq. As expected, P1 cDC1, and P9 cDC1 and cDC2, expressed canonical DC genes *Zbtb46, Flt3, Dpp4*, and *CD24a* (Fig. 2A). Surprisingly, these core genes were also expressed in both P1 XCR1^-^SIRPa^-^ DC and P1 moDC-like cells (Fig. 2A and Fig. S2A). This was especially intriguing with respect to the moDC-like subset as in adult tissues, except in ex vivo contexts (Briseno et al., 2016; Devalaraja et al., 2020), monocyte-derived cells do not express *Zbtb46* and *Flt3* which are generally a feature of precursor-cDC (Pre-cDC) ontogeny rather than monocyte ontogeny (Satpathy et al., 2012). Based on these features we hypothesized these moDC-like cells relate to cDC2s despite CD64 expression. Principal component analysis indicated that DC subsets clustered based on developmental age and type (Fig. 2B). XCR1^-^ SIRPa^-^ DCs transcriptionally aligned with P1 cDC1s, including expression of core cDC1 markers *Itgae*, *Clec9a*, *Irf8,* and *Btla*. Notably, as indicated by our flow cytometric analysis, differential expression of XCR1 was the dominant transcriptional difference between the two populations, along with a small number (11) of other differentially expressed genes.

**Fig. 2.**
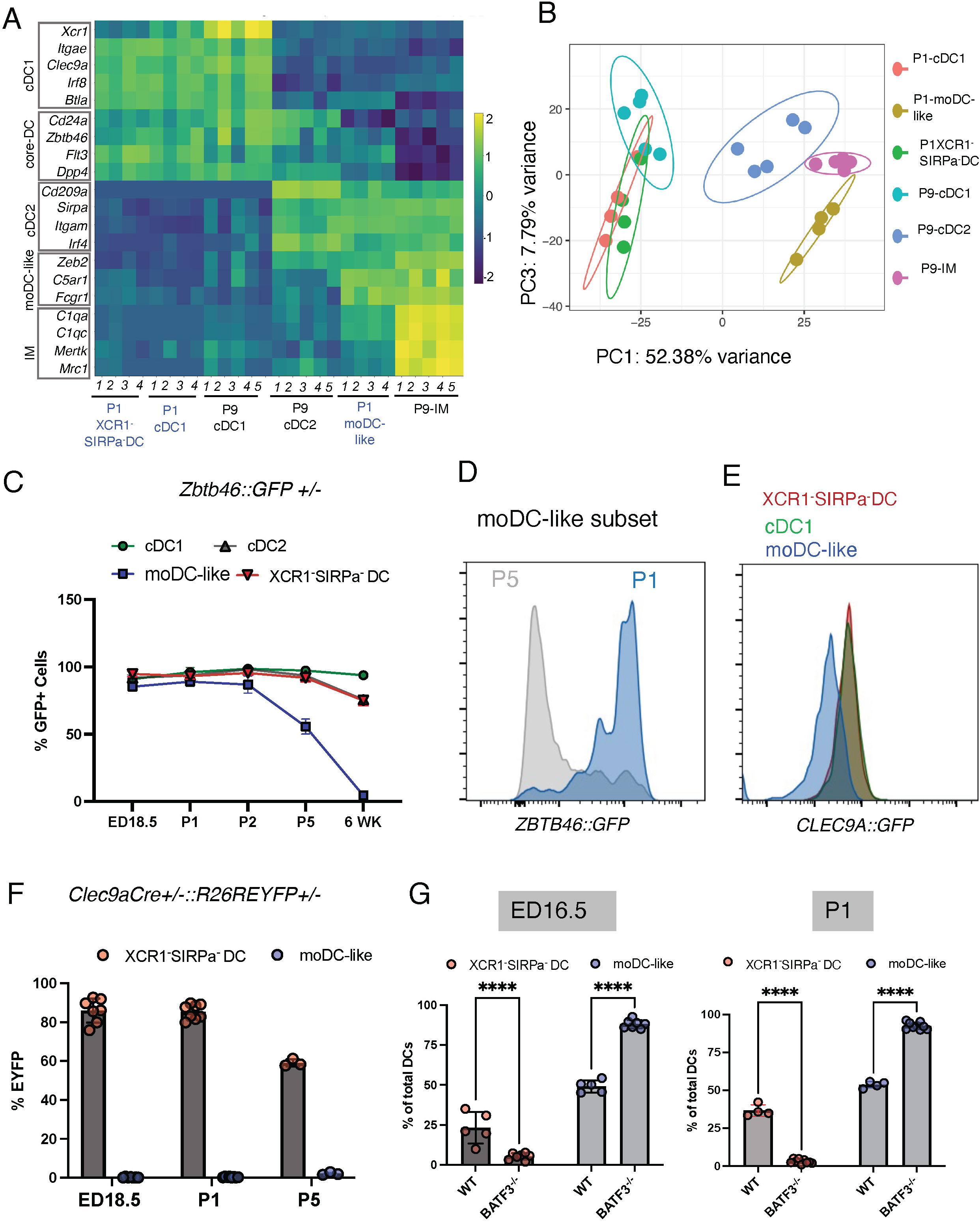
XCR1^-^SIRPa^-^ DCs represent cDC1 in saccular stage lungs. (A and B) Each indicated population was sorted, mRNA isolated, libraries prepared and sequenced. Gene expression heatmap of DEGs scaled by Z score (A), and Principal component analysis (B), among P1 and P9 DC subsets. IMs were used as an out group for contrast. Each replicate represents n=4 mice pooled. (C) Analysis of GFP^+^ cell frequency within each indicated DC subset at specified timepoint in *Zbtb46::GFP^+/-^*mice. (D) Representative histograms showing GFP expression in moDC-like cells at P1 and at P5 time points in *Zbtb46::GFP^+/-^* mice. (E) Representative histograms showing GFP expression in *Clec9a::GFP^+/-^* reporter mice. (F) Bar graph showing percent EYFP+ DC subset at indicated time point in *Clec9aCre^+/-^::R26R-EYFP^+/-^* mice. (G) Frequency of each DC subset at ED16.5 and P1 in WT and *Batf3^-/-^* mice. (C,F,G) Data are shown as mean ± SD. (C-G) Experiments performed two times with n≥5 per group. Significance was determined using 2way ANOVA. ****p < 0.0001.

To further explore the nature of core cDC gene, *Zbtb46*, expression in XCR1^-^SIRPa^-^ DC and moDC-like cells we utilized flow cytometry to analyze GFP expression in *Zbtb46::GFP* reporter mice in each lung DC population across saccular and alveolar developmental stages (Fig. 2C and D). As expected, nearly all cDC1 and cDC2 were positive for GFP expression, at all developmental stages. Consistent with the RNA data, XCR1^-^SIRPa^-^ DCs and moDC-like cells were GFP positive in fetal and early neonatal saccular stage lungs (Fig. 2C). However, the expression of GFP in moDC-like cells waned as lungs transitioned into alveolar stage (P5 to 6 week) suggesting the moDC-like subset may have a developmental origin akin to cDCs.

Transcriptomic analysis clearly identified similarity between the XCR1^-^SIRPa^-^ DCs and XCR1^+^cDC1. To further validate this, we used reporter mice where the cDC-restricted C-type lectin Clec9a promoter drives GFP expression. Prior work has shown that CLEC9A is actively expressed in cDC1 and some pre-cDC in adult mice (Caminschi et al., 2008; Schraml et al., 2013). We first examined expression of GFP in P1 lung DC subsets using *Clec9a::GFP* mice. We found that both cDC1 and XCR1^-^SIRPa^-^DC subsets expressed GFP in contrast to the moDC-like subset (Fig. 2E). Using, a *Clec9a-Cre^+/-^::R26REYFP^+/-^* fate-mapping system, where expression of CLEC9A at any time point of the development allows irreversible marking of the cell with EYFP, we found near complete labeling of XCR1^-^SIRPa^-^DC in early time points as well as cDC1s in adult lungs (Fig. 2F and Fig. S2 C). However, moDC-like cells showed no-labeling at any timepoint. These data suggests that the early life XCR1^-^SIRPa^-^ DCs and moDC-like cells share expression of ZBTB46, suggesting a shared developmental pathway, while these subsets developmentally diverge prior to CLEC9A expression. Next, we tested whether early lung DCs depend on Basic Leucine Zipper ATF-Like Transcription Factor 3 (*Batf3*), a transcription factor necessary for the development of cDC1(Cabeza-Cabrerizo et al., 2021; Hildner et al., 2008). We examined ED16.5 lungs as well as P1 lungs of *Batf3-/-* mice and found that the lungs of these mice were completely devoid of XCR^-^SIRPa^-^ DCs while the moDC-like subset remained unaffected (Fig. 2G and Fig. S2D).

Taken together these data establish that while XCR1^-^SIRPa^-^ DC and moDC-like cells in developing lungs may share a common developmental program (as evidenced by common expression of *Flt3* and *Zbtb46*), they are phenotypically and transcriptomically distinct subsets of DC within newborn lungs. Further, we’ve established that XCR1^-^SIRPa^-^DCs are phenotypically and developmentally equivalent to cDC1 with the exception of a lack of XCR1 expression. From here onwards we propose the term fetal-cDC1 for the XCR1^-^SIRPa^-^DC in these early developmentally immature lungs.

### Cultured ED14.5 MDPs give rise to fetal cDC-like cells

Our data so far establishes the presence of two subsets of DCs, fetal-cDC1s and moDC-like cells, in saccular stage lungs that are phenotypically distinct from alveolar stage lung DCs. While DC development in adults is well characterized (Cabeza-Cabrerizo et al., 2021), whether a similar developmental path exist for early life DCs has not been explored.

In adult mice, DCs originate during bone marrow (BM) hematopoiesis via a series of intermediary progenitors. Among those Macrophage Dendritic Progenitor (MDP) differentiate through Common Dendritic cell progenitor into Pre-cDC, which enter blood circulation, and seed tissues where they further differentiate into tissue cDCs. Development of cDCs is dependent on Fms-like tyrosine kinase 3 ligand (FLT3L)(Ginhoux et al., 2009). Similarly, BM MDPs can also differentiate into monocytes (MO) (Ginhoux and Jung, 2014; Guilliams et al., 2018; Hettinger et al., 2013) and during inflammatory conditions these MO can differentiate into moDCs (Jakubzick et al., 2017). However, BM hematopoiesis only begins towards the end of gestation (Hall et al., 2022), and early myeloid cells such as mast cells, monocytes and macrophages develop from early waves of hematopoiesis that occur in non-bone marrow embryonic sites such as yolk sac and fetal liver (Ng et al., 2023). Thus, we hypothesized that early life DCs have a distinct ontogeny from their adult counterparts.

Fractalkine receptor (CX3CR1) is expressed in yolk sac progenitors, thus studies employing Cx3cr1-Cre fate mapping models have been used to establish macrophage ontogeny in early life. We performed FACS analysis and confirmed that P1 fetal-cDC1 do not express CX3CR1 while moDC-like subset expressed it to a low level (Fig. 3A). This allowed us to use Cx3cr1-Cre::R26R-eyfp fate mapping system to test whether either of the DC subsets derive from a precursor that expressed CX3CR1 in the fetus. FACS analysis of fetal lungs in utero at ED16.5, ED18.5 and at P1 in Cx3cr1-Cre^+/-^::R26R-eyfp^+/-^ mice indicated ∼50% and ∼80% EYFP labeling of Fetal-cDC1 and moDC-like cells respectively (Fig. 3B). This steady labeled cell frequency across fetal to neonatal transition suggests that DC in lungs before birth remain in lungs into early neonatal life. Furthermore, FACS analysis revealed absence of Pre-cDCs (gated as Lin^-^CD45^+^CD11c^+^MHCII^-^SIRPa^-^FLT3^+^) in newborn lungs compared to adults (Fig. S3A). Together these data suggest that newborn fetal-cDC1s and moDCs originate from a non-BM precursor present in the fetal stages that seed lungs prior to birth.

**Fig. 3.**
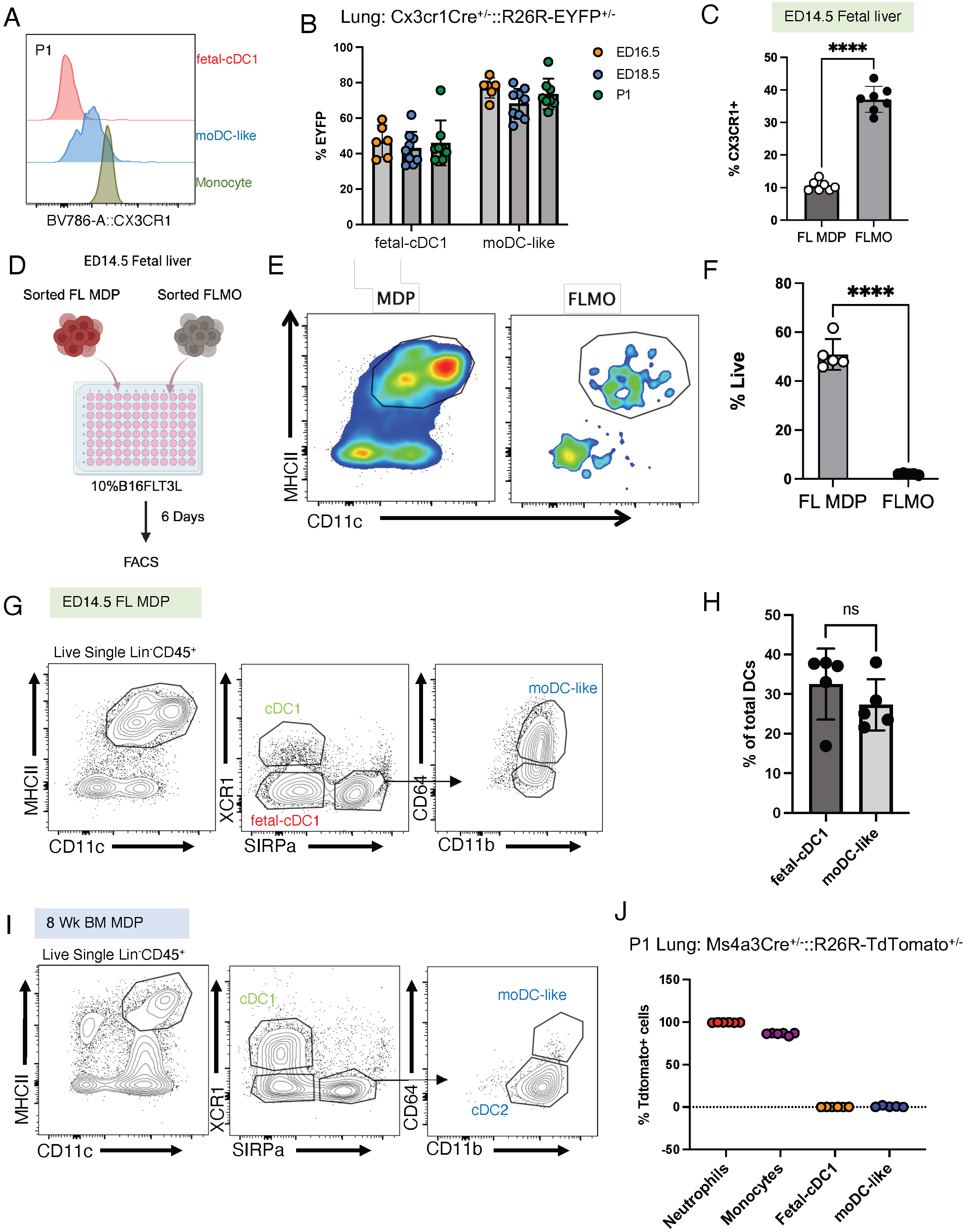
Saccular stage DC subsets derive from fetal precursors. (A) Representative histograms showing CX3CR1 expression on P1 lung DC subsets and monocytes. (B) Percent EYFP positive cells within each lung DC subset, at indicated time point in Cx3cr1Cre^+/-^::R26REYFP^+/-^ mice. (C) Frequency of CX3CR1+ cells within ED14.5 fetal Liver macrophage dendritic progenitors (FL MDP) and FL monocytes (FLMO) analyzed using FACS. (D) Schematic of in vitro culture approach for sorted FL MDP and FLMO from ED14.5 livers. (E) Representative flow plots highlighting CD11c^+^MHCII^+^ cells, 6 days post culture of FL MDP and FLMO in B16FLT3L containing DC media. (F) Bar graph showing percent live cells in FL MDP or the FLMO cultures 6 days post seeding. (G) Representative flow plots showing gating of DC subsets from FL MDP cultures on Day 6. (H) Bar graphs indicate the percent of each DC subset within total DCs in *in vitro* FL MDP cultures. (I) Representative flow plots showing gating of total DC populations (CD11c^+^MHCII^+^) and subsets derived from 8-week old adult bone marrow MDPs (BM MDP) cultured in DC media supplemented with B16FLT3L. (J) Percent TdTomato labeled cells within indicated cell population in Ms4a3Cre^+/-^::R26R-TdTomato^+/-^ mice. A-C and J Experiments were performed two times with n≥4 per group and E-I with n ≥8 pooled fetuses per experiment. Significance was determined using unpaired t test (F), Paired t test (C and H), ****p < 0.0001. ns: not significant.

In the fetus, from ED10.5 to ED15.5 fetal liver becomes the main site of hematopoiesis which harbor highly proliferative early hematopoietic stem cells and their progenitors such as fetal liver(FL) MDP and FLMO (Hoeffel et al., 2015; Hoeffel and Ginhoux, 2018). Adoptive transfers experiments show that FLMO develop into self-renewing alveolar macrophages (Guilliams et al., 2013a; van de Laar et al., 2016) while FL MDP differentiate into monocytes in vitro (Hoeffel et al., 2015) suggesting that FL precursors can give rise to the first myeloid cells in newborn lungs. Therefore, we hypothesized fetal-cDC1 and moDC-like cells in saccular stage lungs originate from fetal precursors, FL MDPs and FLMO respectively. FACS analysis of ED14.5 fetal liver indicated the presence of FL MDP, gated as Lin^-^Sca1^-^MHCII^-^CD45^+^F4/80^-^ CD11c^-^CD11b^-^cKit^+^FLT3^+^ and FLMO gated as Lin^-^Sca1^-^MHCII^-^CD45^+^ F4/80^-^CD11c^-^ckit^-^FLT3^-^ CD11b^+^Ly6C^+^ cells (Fig. S3B), while a subset of FL MDP and FLMO expressed CX3CR1 reflecting previous reports (Fig. 3C) (Hoeffel et al., 2015).

To test whether ED14.5 FL MDP and FLMO develop into DCs, we sorted each subset from ED14.5 fetal livers and cultured them in DC media supplemented with 10% B16FLT3L supernatant (Fig. 3D). As a control we cultured both subsets in culture conditions that received Colony-stimulating factor 1(CSF-1), which drives the differentiation of MDP into MO and macrophages. On day 6 we performed FACS analysis. FL MDP cultured in FLT3L differentiated into DCs while FLMO cultures that received FLT3L died out (Fig. 3E and F). As expected, FL MDP and FLMO that received CSF-1 survived and generated macrophages (Fig. S3C). Strikingly, in FL MDP culture supplemented with FLT3L, distinct populations of CD11c^+^MHCII^+^ cells matching fetal-cDC1 (that lack XCR1 expression) and CD11b^+^CD64^+^ moDC-like cells (Fig. 3G) developed at similar frequency (Fig. 3H).

Next, we tested whether adult BM MDP have similar potential to generate cDCs and moDC-like cells. We isolated BM MDPs and cultured them in FLT3L supplemented media. As expected, BM MDPs develop into cDC1 and cDC2 mirroring previous reports (Fig. 3I), however they did not give rise to CD11b^+^CD64^+^ moDC-like subset indicating that adult BM MDP and ED14.5 FL MDP have distinct potentials. We also cultured adult bone marrow monocytes (BMMo) in FLT3L or CSF-1 and found that BMMo do not survive in FLT3L-containing media while they differentiate into macrophages in CSF-1 supplemented media (Fig. S3 D). Finally, we confirmed that monocytes originating from Granulocyte-monocyte progenitors (GMPs) and their progeny does not give rise to fetal-cDC1 or moDC-like cells, using the recently developed, Ms4a3Cre+/-::R26R-TdTomato fate mapping model which exclusively label GMP-derived cells (Liu et al., 2019). In these mice, FACS analysis for TdTomato labeled cells at P1 indicated near complete labeling in neutrophil and monocyte populations, with no labeling in fetal-cDC1 and moDC-like cells (Fig. 3J and Fig. S3E and F). Taken together these data indicated phenotypically fetal-cDC1 and moDC-like cells develop from ED14.5 fetal liver MDPs that do not go through a monocytic intermediate. Thus, cells previously termed moDC-like in neonatal lungs are in fact ontegenically more similar to cDC2. We thus propose to re-term these cells fetal-cDC2.

### Transcriptome comparisons of fetal-cDC1 and fetal-cDC2 in early lung development

The dendritic cell compartment in early lungs develops against the backdrop of changing environmental conditions accompanied by distinct lung development stages. Furthermore, fetal to neonatal transition at birth has been demonstrated to alter gene expression in a range of cell types (Cohen et al., 2018; Evren et al., 2022; Honda et al., 2022; Li and Magee, 2021; Sun et al., 2022). However, the impact of these environmentally distinct developmental stages on lung DCs has not been explored. To understand how these different developmental environments effect gene expression, we examined the transcriptomes of fetal-cDC1 and fetal-cDC2 from canalicular stage (ED16.5), saccular stage (before birth-ED18.5 and post birth-P1), and early alveolar stage (P5). We performed timed matings to ensure exact developmental stages and harvested lungs at each indicated time point, sorted fetal-cDC1 and fetal-cDC2 based on the gating scheme indicated in Fig. S1C and performed bulk RNAseq.

PCA analysis indicated that fetal-cDC1 and fetal-cDC2 cluster based on the cell type (PC2) over developmental stage (PC1), while similarities between the two DC subsets within a developmental stage were indicated along PC3 (Fig. 4A). Consistent with the PCA results, gene expression analysis clustered fetal-cDC1 and fetal-cDC2 based on cell type rather than the developmental stages (Fig. 4B). Fetal-cDC1 genes similarly expressed across developmental stages include cDC1-specific genes such as *Itgae, Clec9a, Cadm1, Gcsam* and the transcription factor *Irf8* important for cDC1 development(Cabeza-Cabrerizo et al., 2021). fetal-cDC2 across developmental stages (in addition to Zbtb46 and Flt3 as noted above) specifically expressed *Sirpa, Fcgr1* and *C5ar1* (Fig. 4B). In addition, fetal-cDC2 expressed *Cd209* and *Cd14* genes previously shown to express on human and mouse moDCs under inflammatory conditions (Cheong et al., 2010; Marzaioli et al., 2021). Furthermore, across developmental stages, fetal-cDC2s highly expressed several inflammatory cytokine encoding genes compared to fetal-cDC1 (Fig. 4C). These included the myeloid cell chemotaxis associated genes, *Ccl2* and *Ccl9*, neutrophil recruitment associated genes-*Cxcl1* and *Cxcl2*- and acute inflammation associated genes *Tnf* and *Il1b*, suggesting that fetal-cDC2 may particularly play a role in shaping the immune cell milieu in developing lungs.

**Fig. 4.**
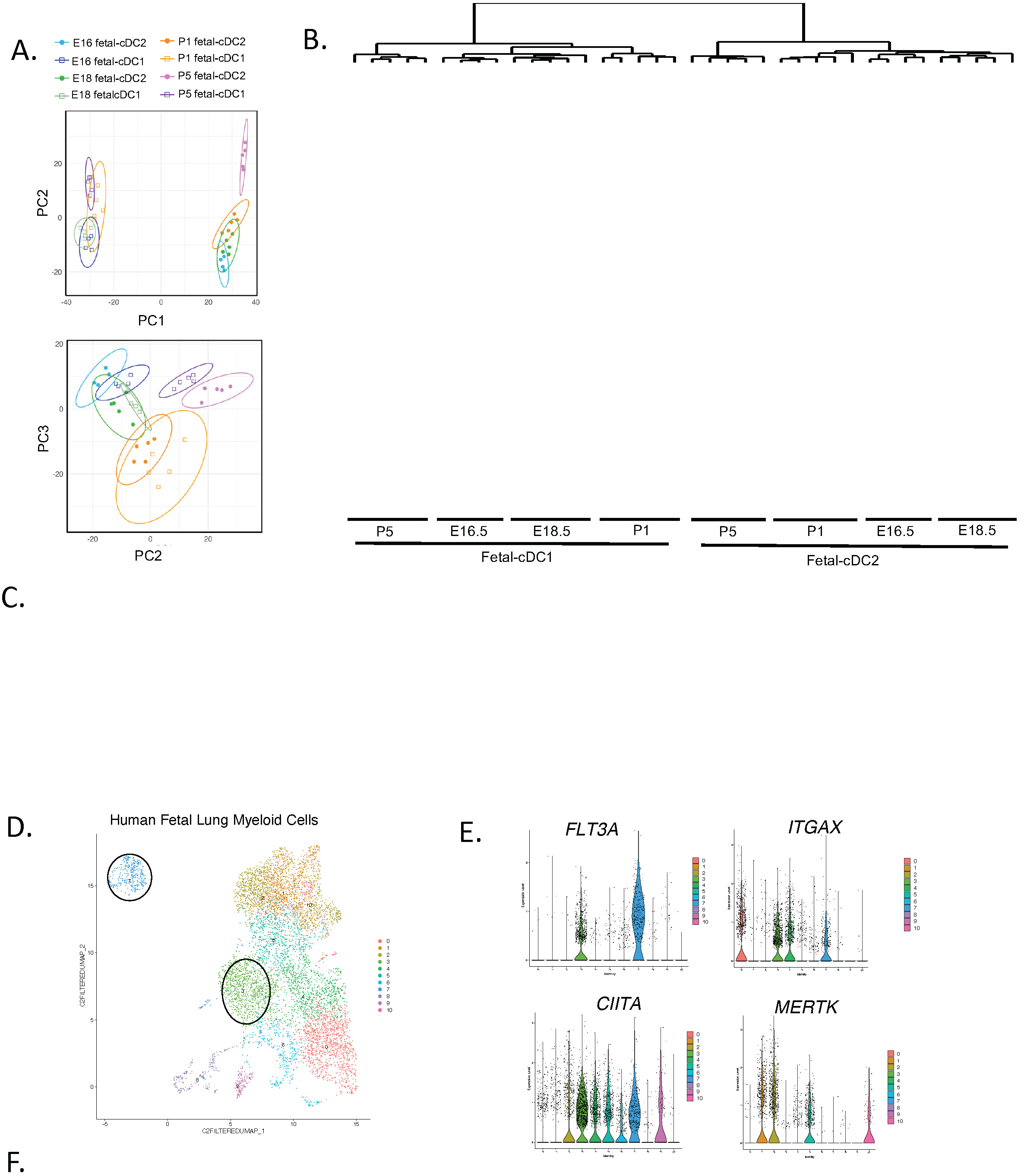
Transcriptome analysis of fetal-cDC1 and fetal-cDC2 across lung developmental stages in mice and human. A-C and D-F are analysis of mouse and human lungs respectively. (A and B) Each indicated population was sorted at the indicated time point from fetal or neonatal mice and analyzed by bulk RNAseq. Principal component analysis (A) and gene expression heatmaps of DEGs scaled by Z score (B). (C) Normalized gene expression of selected inflammatory genes for each DC subset across development. A-C are n=4 or 5 mice per time point. (D) UMAP embedding for human fetal lung single cell RNAseq from publicly available data, highlighting DC clusters 3 and 7 (circled) among other lung myeloid cells. (E) Violin plots showing the expression of DC and macrophage markers on cluster 3 and 7 compared to other myeloid clusters. (F) Violin plots showing DC specific gene expression in cluster 3 and 7.

To investigate whether previously published developing murine lung datasets contained fetal-cDC1s and fetal-cDC2, we visualized the single cell RNAseq (scRNAseq) data set generated by Cohen et. al (Cohen et al., 2018). Our analysis of myeloid cells within this data set established the presence of two clusters (Cluster 14 and 18) that corresponded with *Flt3*, *H2-Ab1* and *Itgax* expression while lacking the macrophage specific gene *Mertk* (Fig. S4B and C). Further analysis revealed Cluster 14 corresponded to fetal-cDC2 with *Sirpa* and *Itgam* expression while Cluster 18 expressed *Clec9a*, *Btla* and *Irf8* corresponding to fetal-cDC1 (Fig. S4 D and E). Cohen. et al data also revealed that cDC1 specific gene *Xcr1* began to express on cDC1-like cells on postnatal day 7 (PND7), while *Clec9a* was expressed from ED16 onwards (Fig. S4 F). Thus, this single cell analysis further validates our findings of XCR1^-^ cDC1, i.e fetal-cDC1 and fetal-cDC2 as the first DCs in developing lungs.

Next, we explored whether DC subsets comparable to fetal-cDC1 and fetal-cDC2 in mice exist in developing human lungs. For this we analyzed scRNAseq data set generated by He et. al. using fetal lungs of 9-22 weeks fetuses (He et al., 2022), which encompass lungs at earlier development stages than the saccular or canalicular stage. In this dataset within the myeloid cells we found two clusters, 3 and 7, that expressed *FLT3A*, and *ITGAX*, but not the macrophage marker *MERTK*, demarcating dendritic cells (Fig. 4 D, E and Fig. S4 G). Further analysis into cluster 3 indicated higher expression of *SIRPA, CX3CR1, CD14, FCGR1A and CD209A* suggesting this cluster represents a fetal-cDC2 subset while cluster 7 with expression of *CLEC9A* and *GCSAM* clearly defined cDC1 (Fig. 4F and Fig. S4H). In contrast to murine fetal-cDC1, these early human fetal lung cDC1s expressed the cDC1 defining marker *XCR1.* Together, transcriptomic data generated here and analysis of public data sets of both human and mice support the presence of unique, stage-specific DC subsets in developing lungs.

### Fetal-cDC1s and *fetal-cDC2* in newborn lungs are functionally mature

Next, we sought to compare the functional maturity of fetal-cDC1 and fetal-cDC2 in early lungs. Specifically with respect to their ability to uptake and process antigens, upregulate MHC Class II molecules and costimulatory molecules such as CD40, CD80 and CD86, engage with naïve T cells and secrete cytokines in order to orchestrate immune responses.

To test their capacity to uptake antigens, process and present to naïve T cells, we sorted each subset from P1 lungs and cocultured them with adult OT-I cells in the presence of OVA peptide. Both fetal-cDC1 and fetal-cDC2 showed equal capacity to induce naïve OT-I cell proliferation (Fig. 5A), indicating that both cells are capable of processing and presenting antigens to CD8 T cells. RNAseq analysis indicated both fetal-cDC1 and fetal-cDC2 in P1 lungs express MHC class II molecules-(*H2-Aa, H2-Ab*) and *Cd40, Cd80, Cd86* costimulatory molecules to a similar level at steady state (Fig. 5B). However, the expression of these genes at protein level (IA/IE, CD40, CD80 and CD86) at baseline were significantly higher on fetal-cDC2 compared to fetal-cDC1 (Fig. 5C) suggesting that at steady state fetal-cDC2 are at an elevated stage of maturation compared to fetal-cDC1. Transcriptome analysis further indicated that early fetal-cDC1 and fetal-cDC2 show differential expression of several Tlr genes (Fig. S5A). *Tlr2*, *Tlr4*, *Tlr7* and *Tlr8* expression were significantly higher on fetal-cDC2 while *Tlr3*, *Tlr11*, and *Tlr12* expression were significantly higher on fetal-cDC1. FACS analysis of P1 fetal-cDC2 and fetal-cDC1 for TLR2 and TLR3 protein expression confirmed the gene expression level differences for these two receptors (Fig. S5B).

**Fig. 5.**
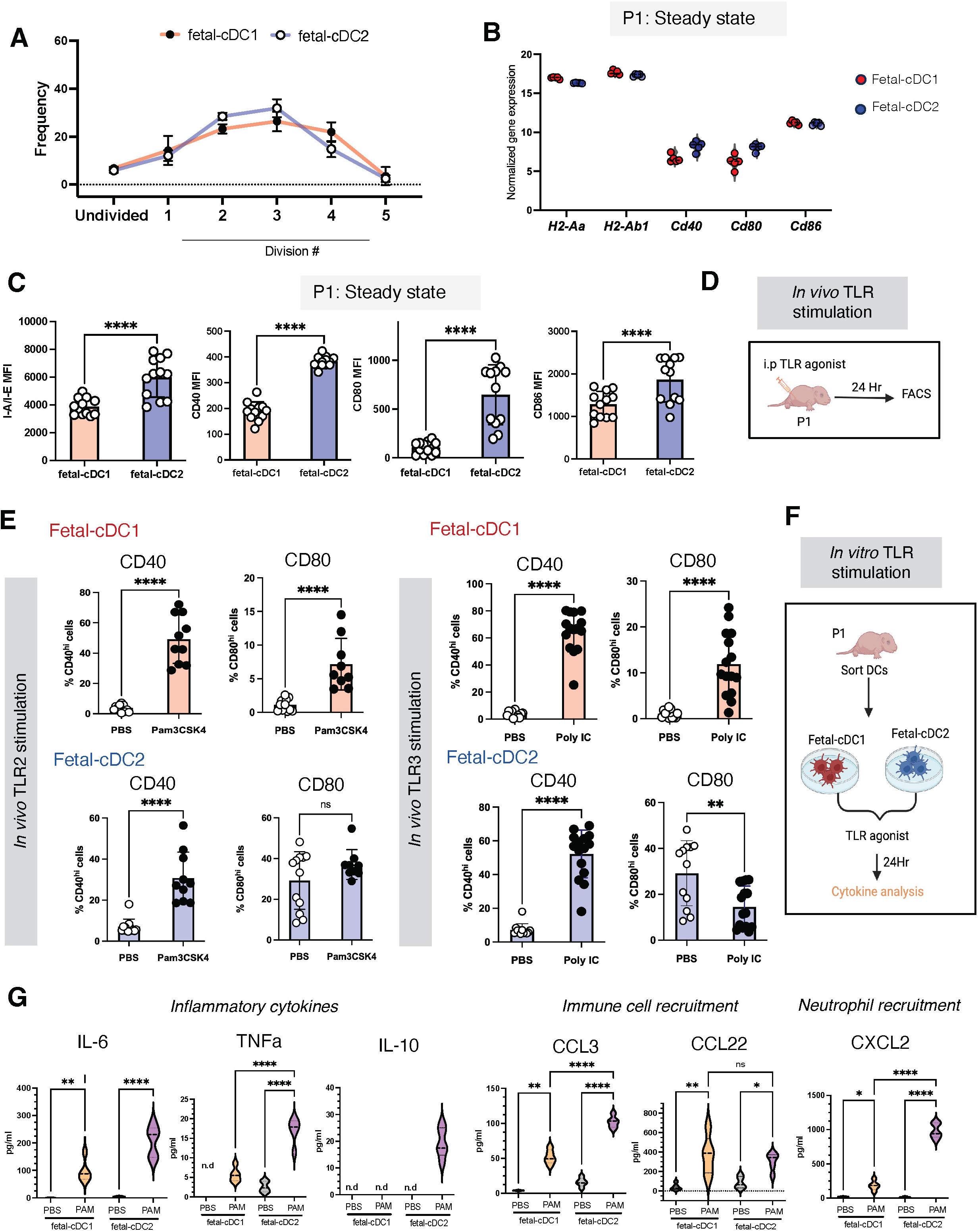
Fetal-cDC1s and Fetal-cDC2s are functionally mature. (A-B) Expression of MHCII and co-stimulatory molecules on P1 fetal-cDC1 and fetal-cDC2 at steady state (A) Normalized gene expression analyzed using RNAseq, n=4-5 per group (B) Mean fluorescent intensity of indicated surface protein, 2 experiments pooled with n≥5 mice per experiment (C) Ovalbumin-specific OT-I CD8+ T cells co-cultured in vitro with primary fetal-cDC1 or fetal-cDC2 sorted from P1 lungs and fed whole OVA protein *ex vivo*. Frequency (mean ± SD) of CFSE+ T cells at each cell division indicated. (D) P1 fetal-cDC1 and fetal-cDC2 were sorted and cultured in the presence of PBS or the TLR2 agonist Pam3CSK4. Levels of indicated inflammatory cytokine analyzed in each supernatant by ELISA at 24 hours post-stimulation. Each violin plot indicates ne=5 samples. n.d: non detected. (E) Percent of each DC subset with CD40 and CD80 positivity identified by FACS analysis on lung DCs post TLR2 or TLR3 agonist administration into P1 neonates. Experiments were performed two times with each condition containing n≥5 neonates per group. B and E Experiments were performed two times with n≥5 per group. C Experiments were performed 2 times with n≥5 pooled P1 lungs per experiment for each of 3 biological replicates. Significance was determined using paired t test (B), unpaired t test (E), one way ANOVA (D), *p < 0.05, **p < 0.01, ***p < 0.001, ****p < 0.0001. ns: not significant.

Next, we interrogated whether P1 DCs respond to TLR2 or TLR3 stimulation in vivo. For this we injected TLR2 or the TLR3 agonists, Pam3CSK or Poly I:C, intraperitoneally to model systemic inflammation (Fig. 5D). Post 24Hr treatment we harvested lungs and performed FACS analysis for activation associated surface protein; CD40 and CD80 in each DC subset. In response to TLR2 and TLR3 stimulation, both fetal-cDC1 and fetal-cDC2 upregulated CD40 and CD80 compared to PBS controls, except in fetal-cDC2 stimulated with TLR3 agonist where CD80 expression lowered compared to PBS (Fig. 5E). These data suggest that despite the baseline differences in TLR2, TLR3, MHCII and coreceptor expression, both subsets are capable of eliciting an activated phenotype to systemic inflammation in vivo.

To test whether this TLR-driven activation of fetal-cDC1 and fetal-cDC2 leads to cytokine production we sorted each subset from P1 lungs, cultured in the presence or absence of the synthetic ligand Pam3CSK4 to stimulate TLR2 or the ligand Poly I:C to stimulate TLR3. After 24 hours of stimulation, we tested the supernatant for 44 different cytokines (Fig. 5F). Post TLR2-stimulation, both DC subsets produced inflammatory cytokines (IL6, TNFa), immune cell recruitment cytokines (CCL3, CCL22) and neutrophil recruitment cytokines (CXCL1, CXCL2) at a significantly higher level compared to the control while anti-inflammatory cytokine IL10 was only detected in the fetal-cDC2 that received Pam3CSK (Fig. 5G). Overall fetal-cDC2 treated with TLR2 agonist indicated a significantly higher level of cytokine secretion compared to fetal-cDC1. Upon TLR3 stimulation, fetal-cDC1 secreted immune cell recruitment cytokine CCL5 to a significantly higher level compared to the control while fetal-cDC2 did not show elevated secretion of any of the cytokines tested compared to the untreated group (Fig. S 5C). Together these in vitro stimulations suggest that both fetal-cDC1 and fetal-cDC2 respond to TLR2 and TLR3 stimulations through cytokine secretion while the type and the quantity of cytokines secreted differ based on the TLR ligand and cell type.

Overall, these *in vivo* and *in vitro* functional studies confirm developing lung DCs are functionally mature and are able to communicate with T cells. While both fetal-cDC1 and fetal-cDC2 respond to TLR-stimulations and produced cytokines, fetal-cDC2 generally secreted increased amounts of cytokines suggesting a more inflammatory phenotype compared to fetal-cDC1.

## Discussion

At birth, fetal lungs transition from in utero aquatic environment to terrestrial air-filled state, signifying not only major changes in mechanical and physiological states, but also in immunological changes. At first breath lungs are exposed for the first time to an array of commensal and pathogenic microbes, and airborne antigens including allergens and pollutants. Thus, newborn lungs need to be equipped with immune cells to intervene, protect and uphold the delicate immune balance. To this end, studies so far have explored the role of DCs in viral and allergy mouse models of lungs spanning the transition from saccular to alveolar stages (Cui et al., 2021; de Kleer et al., 2016; Ruckwardt et al., 2018). However, exploration of DCs in the critical transition of lungs at birth during saccular developmental state has not been undertaken.

Here we explored the DCs in fetal and neonatal mouse lungs and provide the first comprehensive analysis of the DC compartment in saccular development stage of lungs. We found that both major cDC populations (cDC1 and cDC2) exist in developing lungs. We have termed these subsets fetal-cDC1 and fetal-cDC2 as both show distinct phenotypic and ontogenic features from the adult counterparts. The fetal-cDC1s differed phenotypically from adult cDC1 due to the lack of expression of the cDC1-specific surface protein XCR1.

Transcriptomic analysis confirmed the lack of expression of *xcr1* in ED 16.5 to P1 fetal-cDC1. However, except for handful of genes, these fetal-cDC1s transcriptomically resembled a rare population of emerging cDC1s at birth that express intermediate levels of XCR1. A recent study described two distinct cDC1 subsets based on CD103 expression that appears at birth and peak in numbers by day 7, as lungs enter alveolar developmental stage (Silva-Sanchez et al., 2023). The fetal-cDC1s identified here differed phenotypically and kinetically from these CD103+ cDC1 in that they expressed higher level of CD103 during saccular stage and peaked in mid saccular stage by day 2 and waned as lungs developed into alveolar stage. In contrast to adult cDC2, fetal-cDC2 express both *Zbtb46* and *Flt3* in combination with conventionally monocyte-associated markers such as CD64, CD14, and others.

Adoptive transfer experiments (van de Laar et al., 2016) and fate mapping studies have shown that neonatal alveolar macrophages derive from fetal liver monocytes (Hoeffel and Ginhoux, 2018) suggesting that early life myeloid cells are seeded by fetal precursors. However, such analysis for neonatal DCs has not been performed before. Here using in vitro culture systems, we explored the potential of ED14.5 FL MO, FL MDP or adult BM MDP to give rise to phenotypically similar saccular stage lung DCs. Our results indicated that only ED14.5 FL MDP generate both fetal-cDC1 (that lacked XCR1 expression) and fetal-cDC2 that expressed CD64. Consistent with a lack of *Flt3* expression(Hoeffel et al., 2015), ED 14.5 FLMO did not generate DCs, suggesting that FL MDP and FLMO are differentially able to respond to FLT3L cytokine as is required for DC differentiation. Both FLMO and FL MDP responded to CSF-1 and generated macrophages mirroring previous studies. As expected, adult BM MDPs generated cDC1 and cDC2, while a DC subset phenotypically similar to early life fetal-cDC2, that expressed CD64 was absent in this culture system. In animal and human studies moDCs are shown to differentiate from monocytes. However, sorted BMMOs cultured in FLT3L containing media did not give rise to moDCs confirming previous studies that indicated other cytokines such as IL4 and GM CSF are necessary for moDC development (Briseno et al., 2016). An intriguing observation of our study is that only FL MDP showed the capacity to produce phenotypically similar saccular stage lung DCs in response to FLT3L cytokine, suggesting that FL MDP has a higher potential to develop into first lung DCs. Further studies are necessary to determine in vivo capacities of these progenitors to develop into DC subsets.

Our in vitro experiments revealed that both fetal-cDC1 and fetal-cDC2 originate from ED14.5 FL MDP, suggesting a common ontogeny for first lung DCs. This is akin to the adult cDC1 and cDC2 development where both types are generated from a common CDP progenitor and undergo differentiation at the Pre-cDC stage(Cabeza-Cabrerizo et al., 2021). Additionally, we found early life fetal-cDC2 to express ZBTB46 and *Flt3*, defining markers of cDC lineage (Satpathy et al., 2012). Together these lineage data indicate that despite expressing the moDC defining marker CD64, early life moDCs represent the first cDC2s in lungs. This hypothesis parallels human data that show developing fetal lungs contain cDC1 and cDC2 (McGovern et al., 2017). Further studies will be needed to fully elucidate the ontogeny of these cells and whether fetal liver monocytes indeed play no *in vivo* role in lung DC origin.

Functional analysis of fetal-cDC1s and fetal-cDC2 revealed their capacity to communicate with T cells and produce cytokines, suggesting that these first DCs in lungs are in a mature state contrary to previous studies (Ruckwardt et al., 2018). The significantly higher inflammatory gene expression at steady state and increased cytokine production post TLR stimulations of fetal-cDC2 compared to fetal-cDC1 suggest that fetal-cDC2 might be more inflammatory compared to fetal-cDC1 which shows a subdued response in early life. These functional differences, particularly the highly inflammatory nature of fetal-cDC2 may contribute to the exaggerated inflammatory response leading to bronchopulmonary dysplasia, a debilitating condition that can occur in preterm infants, where lungs are at saccular developmental stage.

Previous studies that showed cDC1 are required for early life inflammatory response through IL12 production in viral infections capture the role of DCs in early alveolar stages of lung development (Cui et al., 2023). Further studies that utilize viral or bacterial inflammation models in newborn or in utero are needed to establish whether these first pulmonary DCs play critical roles in shaping lung immune responses to pathogens.

## Materials and methods

### Mice and breeding

C57BL/6J (Jax #000664), B6.129S6(C)-*Zbtb46^tm1.1Kmm^*/J (*Zbtb46^gfp^*, Jax #027618), B6(Cg)-*Clec9a^tm1.1Crs^*/J (*Clec9a^gfp^*, Jax #017696), B6J.B6N(Cg)-*Clec9a^tm2.1(icre)Crs^*/J (*Clec9a^cre^*, Jax # 025523), C57BL/6J-Ms4a3em2(cre)Fgnx/J (*Ms4a3^cre^*, Jax # 036382), B6.129S(C)-*Batf3^tm1Kmm^*/J (*Batf3*^−/−^, Jax #013755), C57BL/6-Tg(*TcraTcrb*)1100Mjb/j (*OT-I*, Jax #003831), B6.Cg-Tg(TcraTcrb)425Cbn/J, B6.Cg-Gt(ROSA)26Sortm14(CAG-tdTomato)Hze/J (TdTomato, Jax #007914) and B6.129X1-*Gt(ROSA)26Sor^tm1(EYFP)Cos^*/J (ROSA-EYFP^fl/fl^, Jax #006148) were purchased from the Jackson Laboratory. Taconic mice (B6-F EF) were purchased from Taconic laboratory. WildR mice were obtained from Oliver J. Harrison at Benaroya Research Institute.

*Zbtb46^gfp^* and *Clec9a^gfp^* male mice were crossed to C57BL/6J female mice to produce heterozygous progeny. *Clec9a^cre^*and *Cx3cr1^cre^* mice were crossed to ROSA-EYFP^fl/fl^ mice to produce heterozygous mice. All mice were bred and maintained in the FHCC vivarium under specific pathogen–free conditions, except for WildR mice, which were bred and maintained in the BRI facility. Male and female mice of each genotype were used. Embryonic development was estimated considering the day of vaginal plug formation as 0.5 days post-coitum (dpc). All fetuses and litters were used in experiments and the minimum sample size used was 3. All procedures involving mice were approved by the FHCC Institutional Animal Care and Use Committee under protocol 51065.

### Tissue isolation and preparation for flow cytometry and sorting

One to 7 days old neonatal mice were euthanized using hypothermia induction followed by decapitation prior to tissue collection. Mice older than 7 days were euthanized using 2.5% Avertin overdose. Pregnant dams were euthanized by CO_2_ overdose followed by cervical dislocation prior to harvesting fetuses. Fetal and neonatal tissue were harvested with the aid of a Leica S9 i microscope. After harvest, fetal and neonatal Lungs, and fetal liver were placed in 3 ml of DMEM (GIBCO) with collagenase-Type IV (0.8 mg/ml, Sigma-Aldrich) and 0.25 mg ml^−1^ DNaseI (Roche). Samples were placed in C-Tubes (Miltenyi), briefly processed with a GentleMACS Dissociator (Miltenyi) and incubated at 37 °C for 30 min and re-processed via GentleMACS. Tissue homogenate was passed through a 100 μm Filter (Fisher). Red blood cells were lysed with 1 ml of 1x RBC lysis buffer (eBioscience) per lung for 5 min at 37 °C and neutralized using FACS buffer. Samples were resuspended in FACS buffer, stained, and analyzed using BD FACSymphony A5 or sorted using BD FACSymphony S6 sorter. Adult lungs were placed in 5ml of DMEM and processed similarly.

### Intraperitoneal administrations

Newborn mice were administered with sterile 1xPBS, 10mg/kg Poly IC (Inviovogen, Poly(I:C) LMW, tlrl-picw) or 10mg/kg Pam3CSK4 (Invivogen, Pam3CSK4:Synthetic triacylated lipopeptide; TLR2/TLR1 agonist, tlrl-pms) in total of 20ul volume intraperitoneally using a Hamilton Syringe (Model 705 RN SYR) and a 34 gauge needle (Hamilton, Small Hub RN Needle, custom length-0.5 inch, point style 4, 12 grade).

### In vitro DC stimulations and cytokine analysis

DCs were sorted and cultured in TC treated 384-well plate (Thermofisher) at 3000 Cells per well. Each well contained 90ul of culture media constituted of DMEM (GIBCO), 10% Fetal Calf bovine serum (GIBCO), 1% L-glutamine (GIBCO), 1% sodium pyruvate (GIBCO), 1% MEM-NEAA (GIBCO), 1% Pencillin-Steptomycin (GIBCO), 55uM 2-mecaptaethanol. At the time of culturing, treatment groups received either 250ng/ml Poly IC (Invivogen) or 50ng/ml Pam3CSK (Invivogen) and the control received PBS. At 24 Hr time point the plate was spun down at 5000 RPM for 6 min at 4C and the supernatant was collected into 0.65 ml snap cap tubes, stored in - 20 C. Collected samples were tested for cytokines by Eve technologies corporation (Calgary, Canada) using Mouse Cytokine/Chemokine 44-Plex Discovery Assay® Array (MD44).

### Fetal liver precursor culture

FACS-sorted ED14.5 fetal liver progenitors were cultured (typically 20000 cells/well) in 96 well flat-bottomed plates. Culture media consisted of 200 μL RPMI 1640 (GIBCO) supplemented with 10% Fetal Calf bovine serum (MiliporeSigma), 1% L-glutamine (GIBCO), 1% sodium pyruvate (GIBCO), 1% MEM-NEAA (GIBCO), 1% Pencillin-Steptomycin (GIBCO), 55 uM 2-mecaptaethanol and 10% B16FLT3L sup or 10ng/ml macrophage colony stimulating factor (CSF-1) (Peprotech). At day 6 cells were harvested on ice, slightly scraping the bottoms of the wells with a pipet tip, passed through a 70 μm filter, washed and stained for flow cytometric analysis.

### In vitro DC-T cell coculture

Sorted DCs were seeded in TC treated 384 well plates 1000 DC per well in 40ul of RPMI media supplemented with 10% fetal Calf serum, 1% L-glutamine, 1% Sodium Piruvate, 1% MEM-NEAA, 1% pencicllin-stepptomycin and 55 uM 2-merceptoethanol. Wells were topped with 40ul of media containing 100ug/ml chicken ovalbumin (OVA) protein (Sigma Aldrich) and 20’000 adult OT-1 cells. T cells were obtained from the spleens of OT-I mice. CD8+ T cells were isolated using EasySep Mouse CD8+ T Cell Isolation Kit (STEMCELL) according to the manufacturer’s protocol. CD8+ T cells were stained with carboxyfluorescein succinimidyl ester (CFSE, 5 μM), washed and T cells were cultured with DCs at 20:1 ratio for 3 days. T cell proliferation was assessed by CFSE dilution using flow cytometry.

### RNAseq sample preparation and data analysis

Fetal and neonatal Lung cells were stained for flow cytometry and DC populations sorted on BD FACSymphony S6 sorter. Sorted cells (200 per population) were immediately processed using SMART-Seq™ v4 Ultra™ Low Input RNA Kit for Sequencing (Clonetec laboratories) per manufacturer’s instructions. The amplified library was run on a Nextseq 2000 at the Benaroya Research Institute Genomics Core. The resulting data was processed using ROSALIND® (https://rosalind.bio/)software, with a HyperScale architecture developed by ROSALIND, Inc. (San Diego, CA). Reads were trimmed using cutadapt (Martin, 2011). Quality scores were assessed using FastQC (https://www.bioinformatics.babraham.ac.uk/projects/fastqc/). Reads were aligned to the Mus musculus genome build mm10 using STAR (Dobin et al., 2013).

Individual sample reads were quantified using HTseq (Anders et al., 2015) and normalized via Relative Log Expression (RLE) using DESeq2 R library (Love et al., 2014). Quality control metrics were generated using RSeQC (Wang et al., 2012). DEseq2 was used to calculate fold changes and p-values for differentially expressed genes. Heatmaps and additional analysis were performed in R and R studio using DEseq2 and Plotly for data visualization.

### Cell clustering and DEG profile

Public single cell RNA seq data was download from NCBI Geo. Murine data (Cohen et al., 2018) accession # GSE119228. Human data (He et al., 2022) were downloaded from CellxGene. Data were imported into Seurat. Standard Seurat workflows were used to perform UMAP based dimensionality reduction, cell clustering, cell type identification, and DEG.

### Imaging

Newborn or adult mice were euthanized per protocol and lungs were harvested into containers pre-filled with 10% neutral buffered formalin. Tissues were allowed to fix for 3 days in room temperature and processed for imaging by the Fred Hutch Experimental Histopathology core.

### Quantification and Statistical analysis

Data were analyzed with Prism 10 (GraphPad Software). Statistical details and group sizes are indicated in the figure legends.

## Data availability

Sequencing results are available from the NCBI Geo under accession number (TBD). All other data are available in the main text or the supplementary materials. All experimental models and reagents will be made available upon installment of a material transfer agreement. Further information and requests for resources and reagents should be directed to and will be fulfilled by the lead contact (M.B. Headley).

## Supplemental material

This article includes five supplemental files that show flow cytometry gating strategies, representative plots, and supplemental experimental figures. Fig. S1, relating to Figs. 1 show Phenotypical analysis of early life Dendritic cells. Fig. S2, relating to Figs. 2 show XCR1-SIRPa-DCs resemble cDC1. Fig. S3, relating to Fig. 3 Shows Fetal Liver precursors give rise to first DCs in lungs. Fig. S4, relating to Fig. 4 shows Transcriptome analysis of DC subsets in murine and human developing lungs. Fig. S5, relating to Fig. 5, Functional analysis of fetal-cDc1 and moDC

## Acknowledgements

We thank Fred Hutch Cancer Center Flow Cytometry Core, Histopathology Core, Comparative Medicine, and the Genomics core at the Benaroya Research Institute for their support.

The study was supported by a Hartwell foundation grant and NIH R21 AI173825 to Mark. B Headley.

We would also like to thank Dr. Oliver Harrison for access to WildR mice. As well as Dr. Joshua Pollack for assistance with bulk RNAsequencing analysis.

## Author contributions

R. Soysa: Conceptualization, Data curation, Formal analysis, Investigation, Methodology, Software, Validation, Visualization, Writing - original draft, Writing - review & editing; S. Abideen: Data curation, Methodology; V. Zepeda Reyes: Methodology and Animal study support. M.B. Headley: Conceptualization, Data curation, Formal analysis, Software, Funding acquisition, Resources, Supervision, Writing - review & editing.

---

**Fig. S1.**
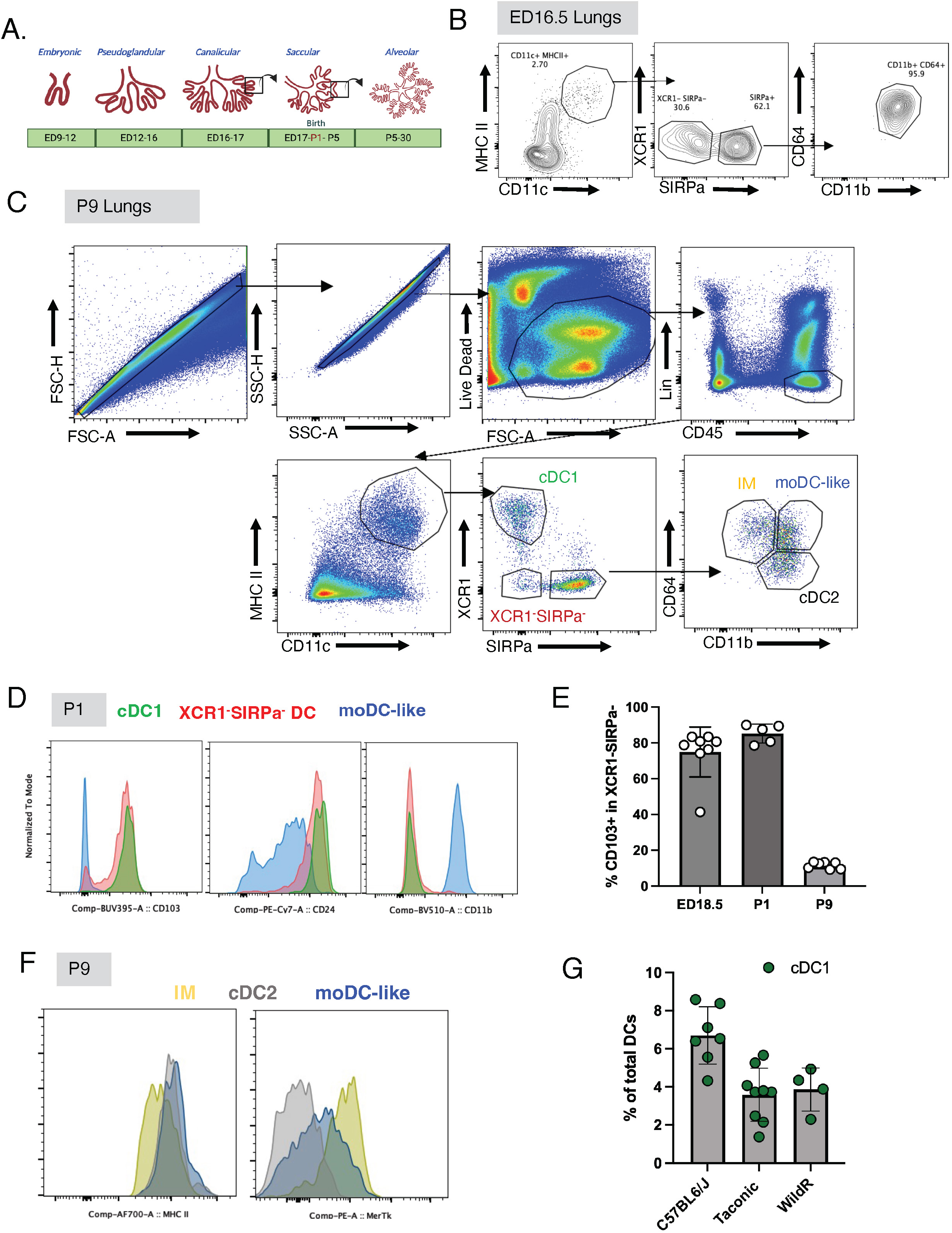
Phenotypical analysis of early life Dendritic cells. (A)Schematic of lung developmental stages in mice. ED: Embryonic Day, P: Postnatal (B) Representative FACS plots indicating DC subsets in ED16.5 lungs (C) Representative FACS plots indicating the complete gating strategy to identify neonatal DC subsets. cDC: conventional-DC, moDC-like: Monocyte-derived DC-like, XCR1^-^SIRPa^-^DC and IM: interstitial macrophages. Lineage:CD3 B220 CD19 NK1.1 SiglecF NKp46 Ter119 (D)Representative histograms indicating expression of CD103, CD24 and CD11b on P1 DC subsets. (E)Percent CD103^+^ DCs in XCR1^-^SIRPa^-^DCs at indicated time point. (F) Representative histograms showing surface expression of MERTK and MHCII in IMs, moDC-like cells and cDC2 in P9 lungs. (G) Bar graphs showing cDC1 frequency within total DCs in 3 different colonies of C57BL6 mice. B-G experiments were performed 2 times with n=4 or more mice per group.

**Fig. S2.**
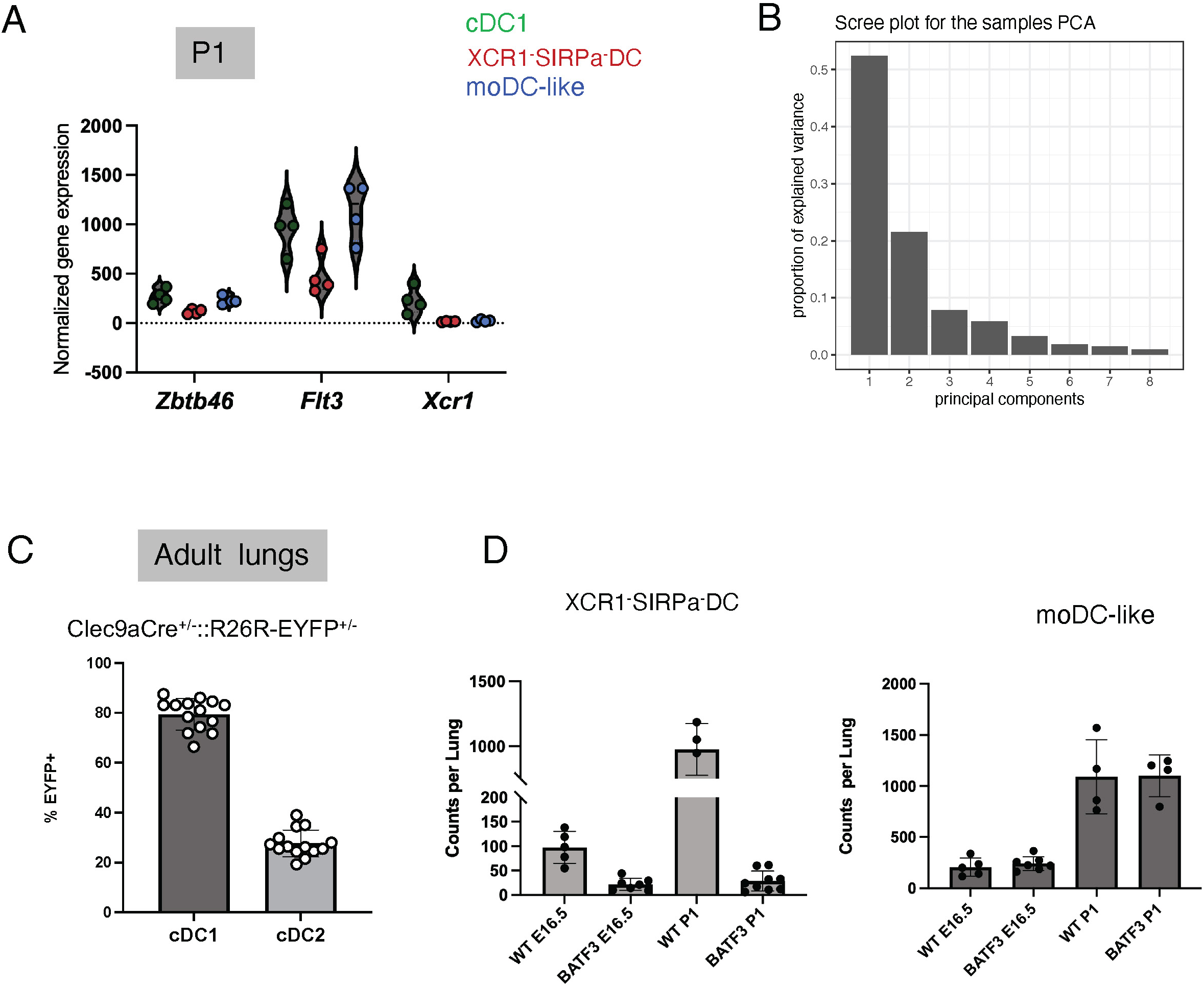
XCR1^-^SIRPa^-^ DCs closely resemble cDC1. (A) Canonical DC genes, *Xcr1*, *Flt3* and *Zbtb46* expression on P1 DC subsets determined using RNAseq analysis. Each replicate represents n=4 mice pooled (B) Scree plot showing the variance of each principal component relating to RNAseq data represented in Fig 2B. (C) Bar graph showing percent EYFP^+^ cDC1 and cDC2 in lungs of adult Clec9aCre^+/-^::R26REYFP^+/-^ mice. (D) DC subsets plotted as counts per lung at ED16.5 and P1 in WT and Batf3^-/-^ mice (related to Fig 2G) Data are shown as mean ± SD. C-D experiments performed two times with n≥5 per group.

**Fig. S3.**
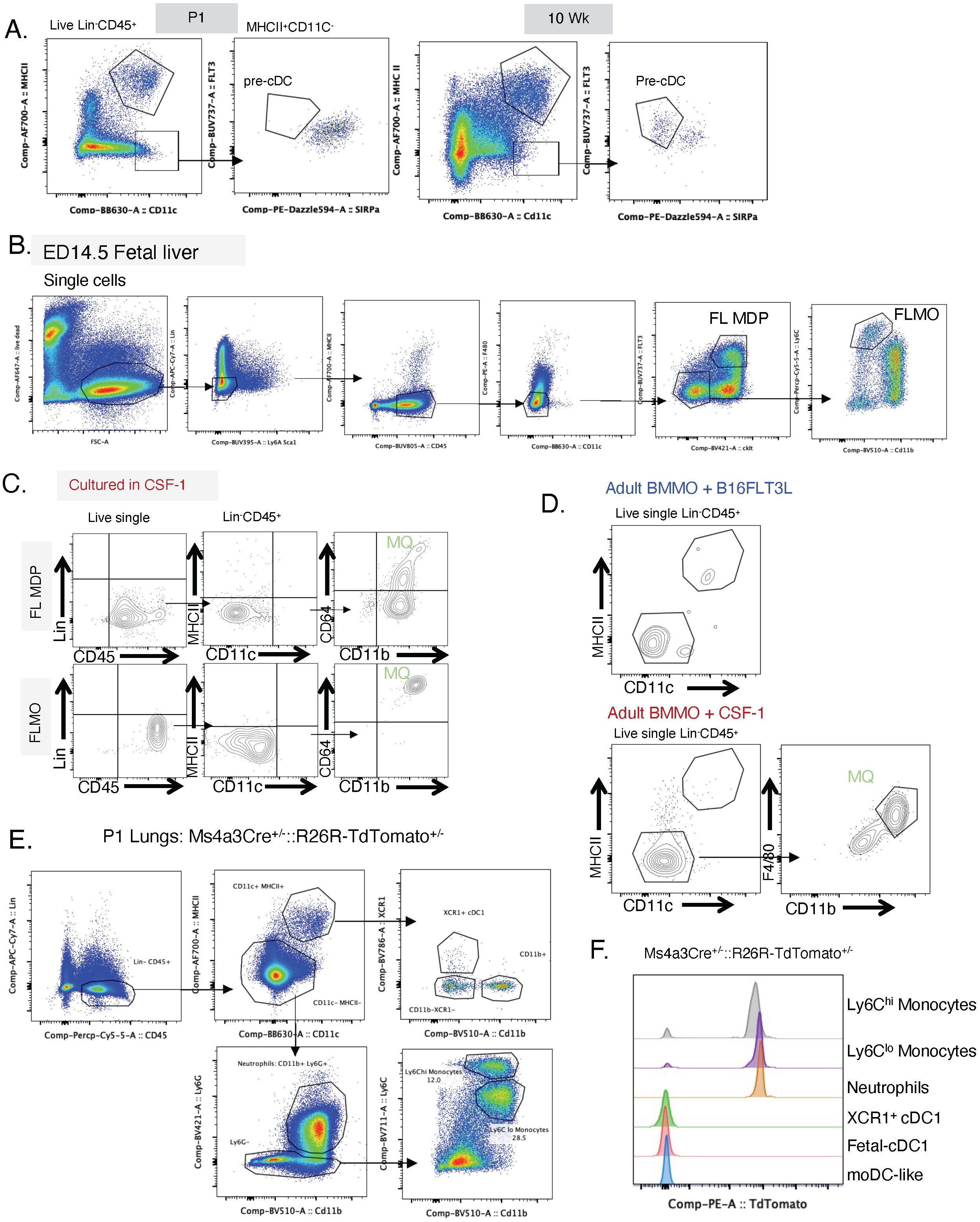
Fetal Liver precursors give rise to first DCs in lungs. (A) Representative FACS plots highlighting gating of pre-cDC in P1 and 8-week adult lungs within live cells. Pre-cDCs in lungs were gated as Lin^-^ CD45^+^ CD11c^+^ MHCII^-^ SIRPa^-^ FLT3^+^. (B) Representative FACS plots indicating gating for ED14.5 fetal liver (FL) MDP and FLMO. (C) Representative FACs plots indicating FL MDP and FLMO cultured in CSF-1 containing media for 6 days. MQ: macrophages. (D) Representative FACs plots showing DC and macrophage populations in adult BM MO cultured in B16FLT3L or CSF-1 containing media for 6 days. (E) Representative FACS plots of gating myeloid subsets and (F) representative histograms indicating TdTomato labeling in Ms4a3Cre^+/-^::R26R-TdTomatao^+/-^ mice at P1. A-F experiments are performed twice. A,B,E,F with n≥4 per group. C with n≥8 fetuses pooled and D with n≥2 adult mice pooled.

**Fig. S4.**
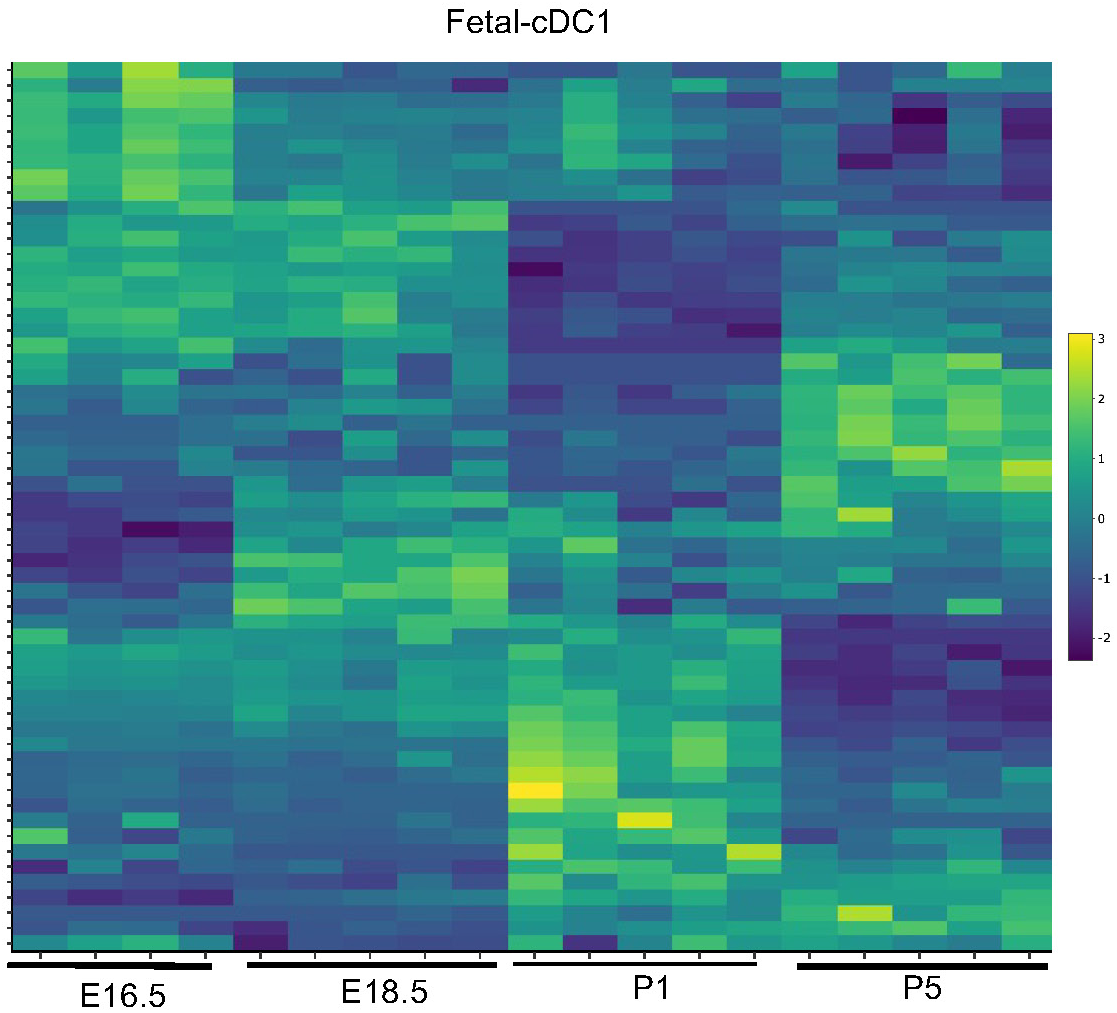
Transcriptome analysis of DC subsets in murine and human developing lungs. A-F and G-H represent mice and human respectively. (A) Gene expression analyzed using bulk-RNAseq for each DC subset at the indicated time point visualized as a heatmap, scaled by Z score. (B) UMAP derived from scRNAseq data produced by Cohen. et al for developing murine lungs highlighting DC clusters 14, relating to cDC1 and cluster 18 relating to fetal-cDC2. (C) Expression of indicated gene in the DC clusters 14 and 18 in panel B. (D and E) Violin plots showing fetal-cDC1 and fetal-cDC2 specific gene expression in clusters 14 and 18. (F) Violin plots showing cDC1 genes, *Clec9a* and *Xcr1* expression at the indicated time point for cluster 18, cDC1s. (G) UMAP showing the emergence of DC subsets in human fetal lungs from 9-22 weeks of gestation derived from He at. al. scRNAseq data. (H) Fetal-cDC2 specific genes *FCGR1A* and *CD209A* expression in clusters 3 and 7 related to Fig 4D.

**Fig. S5.**
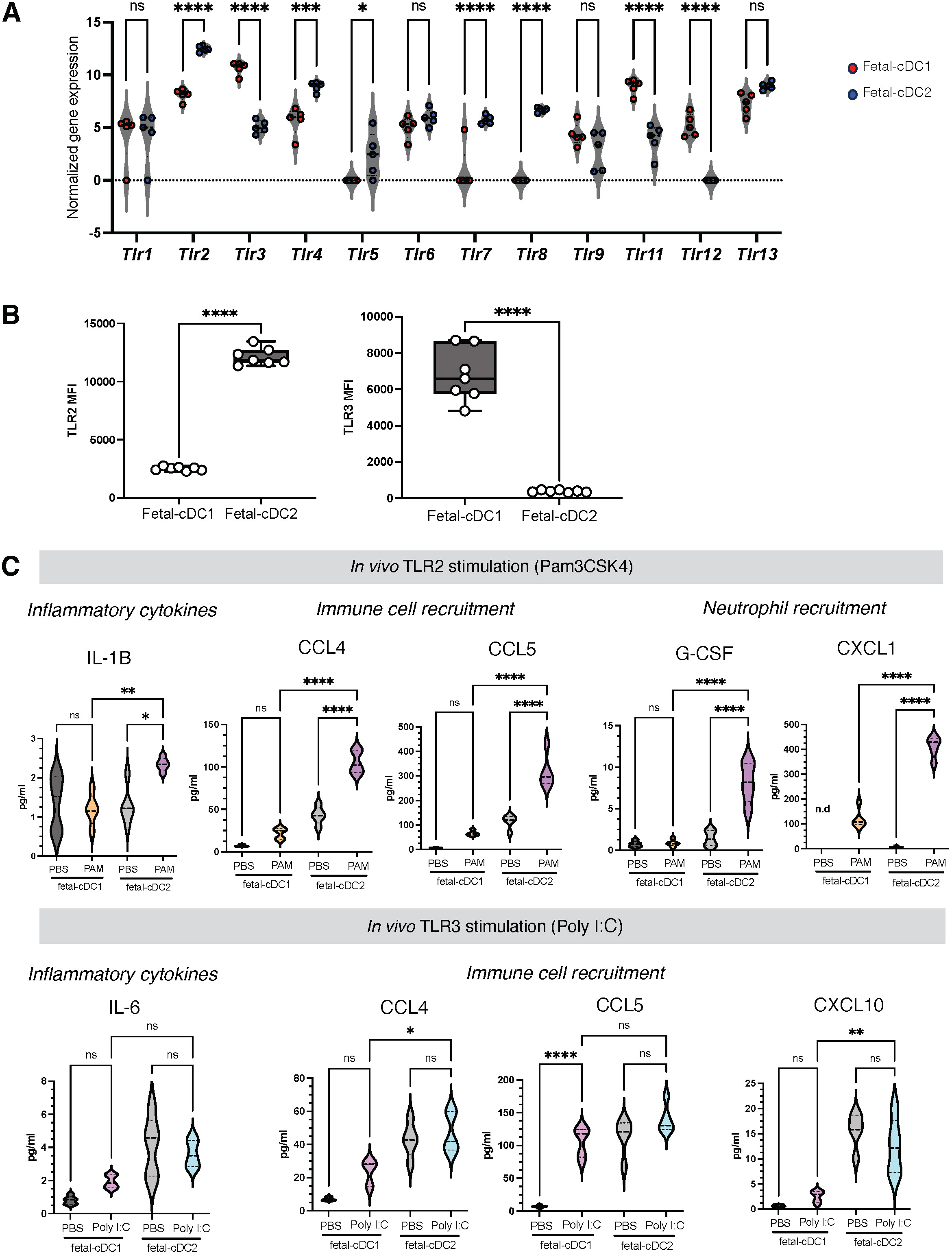
Functional analysis of fetal-cDc1 and moDC. (A) Normalized gene expression of *Tlr* receptors from RNAseq for each DC subset in P1 lungs. Each plot represents n=5 samples. (B) Mean fluorescence intensity of TLR2 and TLR3 surface expression on each DC subset in P1 mouse lungs. Data indicate Two experiments pooled n≥5 per experiment. (C) Levels of indicated cytokine analyzed in each supernatant, by ELISA, 24 hours post culture of each sorted P1 DC subset with Pam3CSK or Poly I:C. Each violin plot represents n=5 samples. n.d: non detect. Significance was determined using 2-way ANOVA (A), paired t test (B), one way ANOVA (D), *p < 0.05, **p < 0.01, ***p < 0.001, ****p < 0.0001. ns: not significant.

## Notes

### Competing Interest Statement

The authors have declared no competing interest.

